# GCBM-DCT-HV-Bio-NL-Grow-CHG-CSM-RHEC: A Unified Geometric, Biological, Causal, and Regenerative Framework for Mechanism-Aware Tissue and Connectome Modeling

**DOI:** 10.64898/2026.06.24.734320

**Authors:** Tao Xu, Zhixin Hu, Xiaodian Sun, Li Jin, Momiao Xiong

## Abstract

Modern biological prediction problems increasingly require models that go beyond Euclidean feature regression and local graph smoothing. Tissue, cellular, and connectome systems are nonlinear, geometry-dependent, intervention-sensitive, history-dependent, and subject to regenerative or homeostatic constraints. We propose **GCBM-DCT-HV-Bio-NL-Grow-CHG-CSM-RHEC**, a unified model for mechanism-aware biological prediction. The model integrates geometric connectome dynamics, differentiable charted tissue geometry, Hamiltonian latent transport, nonlinear biological kinetics, nested latent memory, continual growth without overwriting, causal hypergraph structure, causal structure modeling, and regenerative homeostatic error correction. Unlike Euclidean baselines, which treat observations as flat vectors, and local graph baselines, which use neighborhood smoothing without mechanistic structure, the proposed model represents biological states (Trapnell 2015) as coupled geometric, dynamical, causal, and regenerative objects.

We evaluate the model on four synthetic toy studies, Toy A–D, designed to reflect increasing biological complexity: local Euclidean structure, nonlinear mechano-chemical interaction, causal intervention response, and out-of-distribution regenerative shift. Compared with Euclidean and local graph baselines, the full model achieves the lowest mean squared error across all four toy studies. Relative to the Euclidean baseline, the full model reduces MSE by approximately 63.0%, 89.1%, 89.0%, and 90.9% on Toy A, Toy B, Toy C, and Toy D, respectively. These results support the value of integrating geometry, mechanism, causal structure, adaptive growth, and regenerative correction into a single predictive architecture (Figure 1).

## 1. Introduction

Current omics analysis and drug-development pipelines often rely on simplified representations of biological systems. Gene expression, mutation status, molecular fingerprints, and pathway annotations are frequently treated as high-dimensional but essentially Euclidean feature vectors. Although this abstraction has enabled large-scale modeling, it often neglects key determinants of biological function, including cell and tissue morphology, mechanical forces, spatial organization, causal intervention structure, temporal history, microenvironmental feedback, and regenerative or homeostatic constraints.

Over the past several decades, many computational benchmarks and synthetic biological datasets have been generated under simplified assumptions: linearity, low-dimensional Euclidean structure, weak or absent mechano-chemical coupling, stationary distributions (Kim et al. 2026), and limited causal or tissue-level context. As a result, models trained and evaluated on such data may appear accurate under idealized conditions while failing to capture the complexity of real biological systems. That mismatch can make benchmarks look successful while failing to translate to real biological or clinical impact. This mismatch may contribute to poor translation from preclinical discovery to therapeutic (Zhnag t al. 2026) efficacy, where drug candidates often fail because cellular, tissue, organism-level, or context-specific responses were not adequately represented.

A more realistic direction for biological research is to move beyond flat omics-only prediction toward mechanism-aware, multi-modal, and multi-scale modeling (Song and Zhen 2026). Future predictive systems should integrate omics, morphology, tissue geometry, mechanical forces, perturbation history, causal structure, cell-cell interaction, and regenerative or homeostatic feedback. Such integration would better reflect how biological systems actually respond to drugs, genetic perturbations, injury, aging, and disease. The goal is not merely to build more complex models, but to align model structure with biological reality.

There is also real support for this direction. Drug development remains highly attritional; one analysis notes high attrition and limited annual approvals in clinical development, and another JAMA Internal Medicine study found that roughly half of investigational drugs entering late-stage development fail during or after pivotal trials, mainly because of safety, efficacy, or both (Zhou .et al. 2025; Hwang et al. 2016). Mechanobiology research emphasizes that mechanical forces shape cellular behavior, tissue architecture, and disease progression, while Cell Painting (Seal et al. 2025) and image-based profiling studies show that morphology can capture biologically meaningful perturbation responses and support drug-discovery applications (Kaluk et al. 2025; Seal et al. 2025).

Biological AI should not only predict from what is easy to measure; it should help redesign what we measure. To improve drug discovery and biological prediction, we need datasets and models that include the physical, spatial, causal, temporal, and regenerative structure of living systems. Biological AI should not only predict from what is easy to measure; it should help redesign what we measure. To improve drug discovery and biological prediction, we need datasets and models that include the physical, spatial, causal, temporal, and regenerative structure of living systems. To address these limitations, we propose **GCBM-DCT-HV-Bio-NL-Grow-CHG-CSM-RHEC**, a full unified framework (Wang et al. 2026; Zhang et al 2026; Zhang 2026) for biological prediction and mechanism discovery. The model is designed as a complete architecture, not as an incremental extension of a previous partial model. Its purpose is to jointly represent:

1. geometric connectome flow (Zhang 2026);
2. differentiable tissue charts (Xie et al. 2026);
3. Hamiltonian latent transport and verification (Schiebinger 2019; Zhang 2026) ;
4. nonlinear biological kinetics (Evans et al. 2025);
5. nested latent and memory structure (Biswas et al. 2026);
6. adaptive growth without overwriting (Adila et al. 2026; Li et al. 2019);
7. causal hypergraph interactions (Harit and Sun 2025);
8. causal structure modeling (Ejaz and Bareinboim 2026; Orujlu et al. 2026);
9. regenerative homeostatic error correction (Richter et al. 2026; Mohan, and Surulescu 2025).

The central hypothesis is that biological prediction improves when the model jointly learns **where the system is, how it moves, which mechanisms are active, which interventions are causal, which structures should be preserved**, and **how the system restores or regenerates toward biologically plausible states**.

## 2. Model Overview

Let each biological sample or time point be represented by

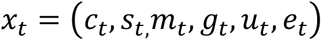

where *c*_*t*_ denotes chemical or molecular signals, *s*_*t*_ denotes mechanical stress or stiffness, *m*_*t*_ denotes tissue morphology, *g*_*t*_ denotes graph or connectome structure, *u*_*t*_ denotes intervention variables, and *e*_*t*_ denotes environmental or experimental context. The goal is to predict a biological response

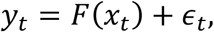

where *F* may be nonlinear, geometry-dependent, intervention-sensitive, and mechanism-dependent.

The proposed model learns a structured latent state

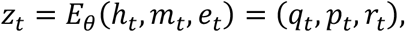

where *q*_*t*_ represents configuration-like biological state, *p*_*t*_ represents momentum-like direction of transformation, and *r*_*t*_ represents causal mechanism coordinates. The final prediction is obtained from a mechanism-aware decoder:

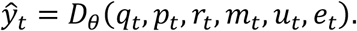

The model is trained by minimizing a composite objective:

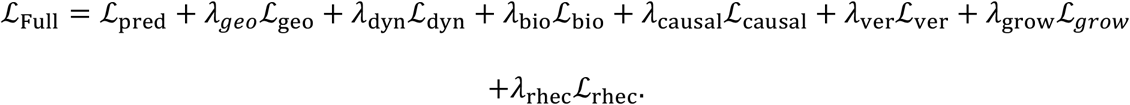

The prediction loss is

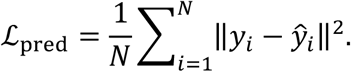

The remaining terms enforce geometry preservation, dynamical consistency, biological mechanism consistency, causal structure, verification (Wan et al. 2026), continual growth (Hassani et al. 2026), and regenerative homeostatic correction.

## 3. Methods

### 3.1 Baseline Models

We compare the full model with two baselines.

#### 3.1.1 Euclidean Baseline

The Euclidean baseline treats all observations as flat vectors:

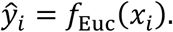

It does not encode graph structure, manifold geometry, causal intervention structure, latent biological mechanisms, or regenerative correction. It is included to test whether the response can be explained by raw covariates alone.

#### 3.1.2 Local Graph Baseline

The local graph baseline incorporates neighborhood information through local smoothing or graph aggregation:

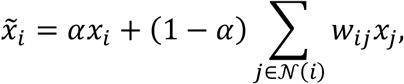

followed by a predictor

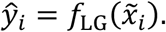

This baseline tests whether local graph context improves prediction. However, it does not distinguish causal from non-causal neighbors, does not model nonlinear biological mechanisms, and does not include regenerative or homeostatic constraints.

The full model differs from both baselines in the components described below.

### 3.2 GCBM: Geometric Connectome Biological Model

The **GCBM** component represents biological activity as structured flow over a signed connectome or tissue interaction graph. Let

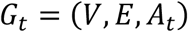

be a time-dependent graph, where *V* denotes biological units such as cells, tissue regions, or connectome nodes; *E* denotes interactions; and *A*_*t*_ is a weighted, possibly signed adjacency matrix.

Each node has a hidden state *h*_*i,t*_. The connectome flow is modeled by

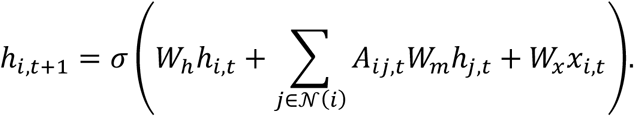

Unlike local graph smoothing, GCBM does not simply average neighboring features. It learns signed and directed biological message flow (Yang et al. 2019), allowing excitatory, inhibitory, cooperative, and antagonistic interactions.

The connectome state is

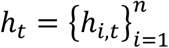

This state becomes the input to the geometric, latent, causal, and regenerative modules (Figure 2).

**Figure 1.**
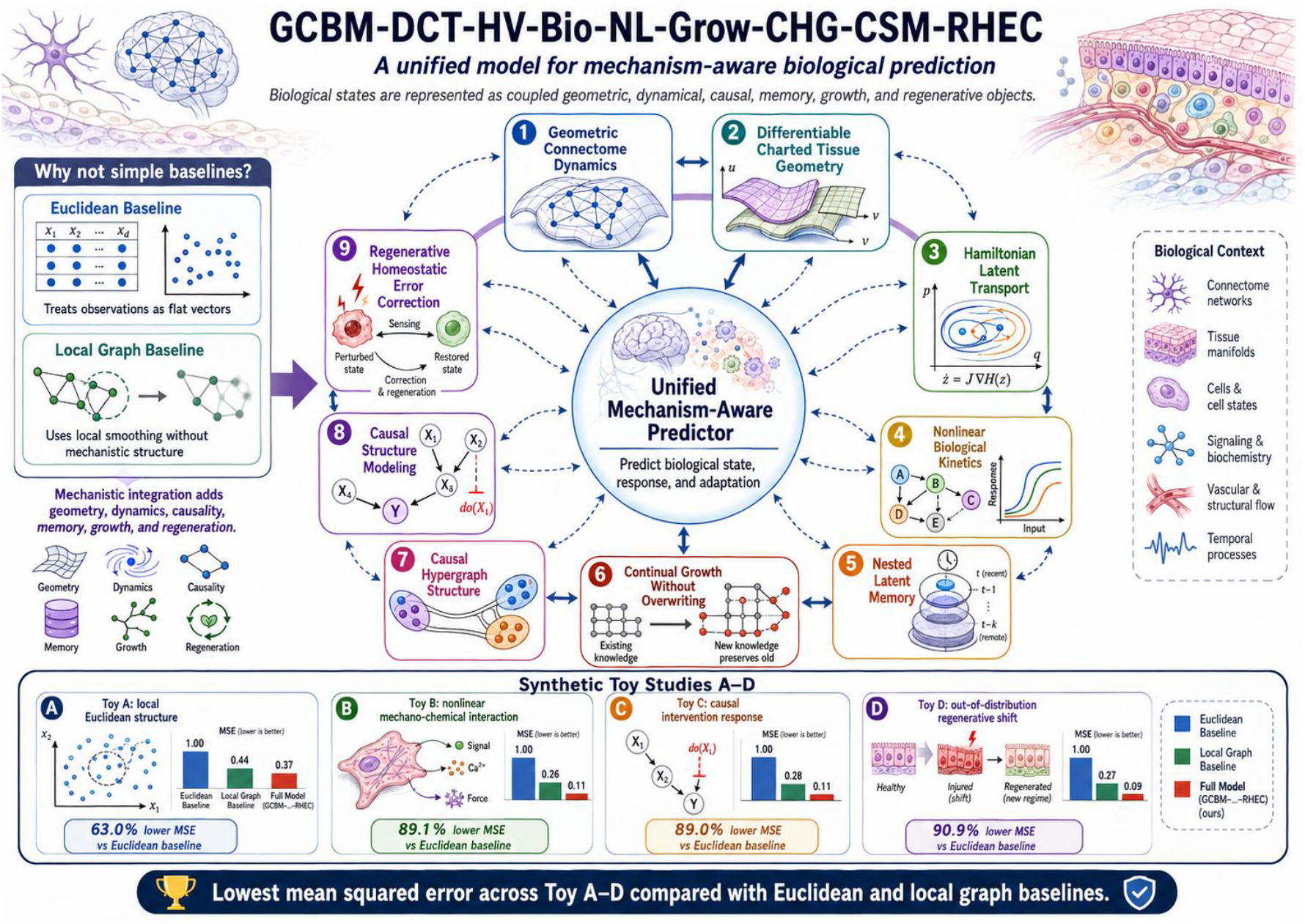
GCBM-DCT-HV-Bio-NL-Grow-CHG-CSM-RHEC, a unified model for mechanism-aware biological prediction.

**Figure 2.**
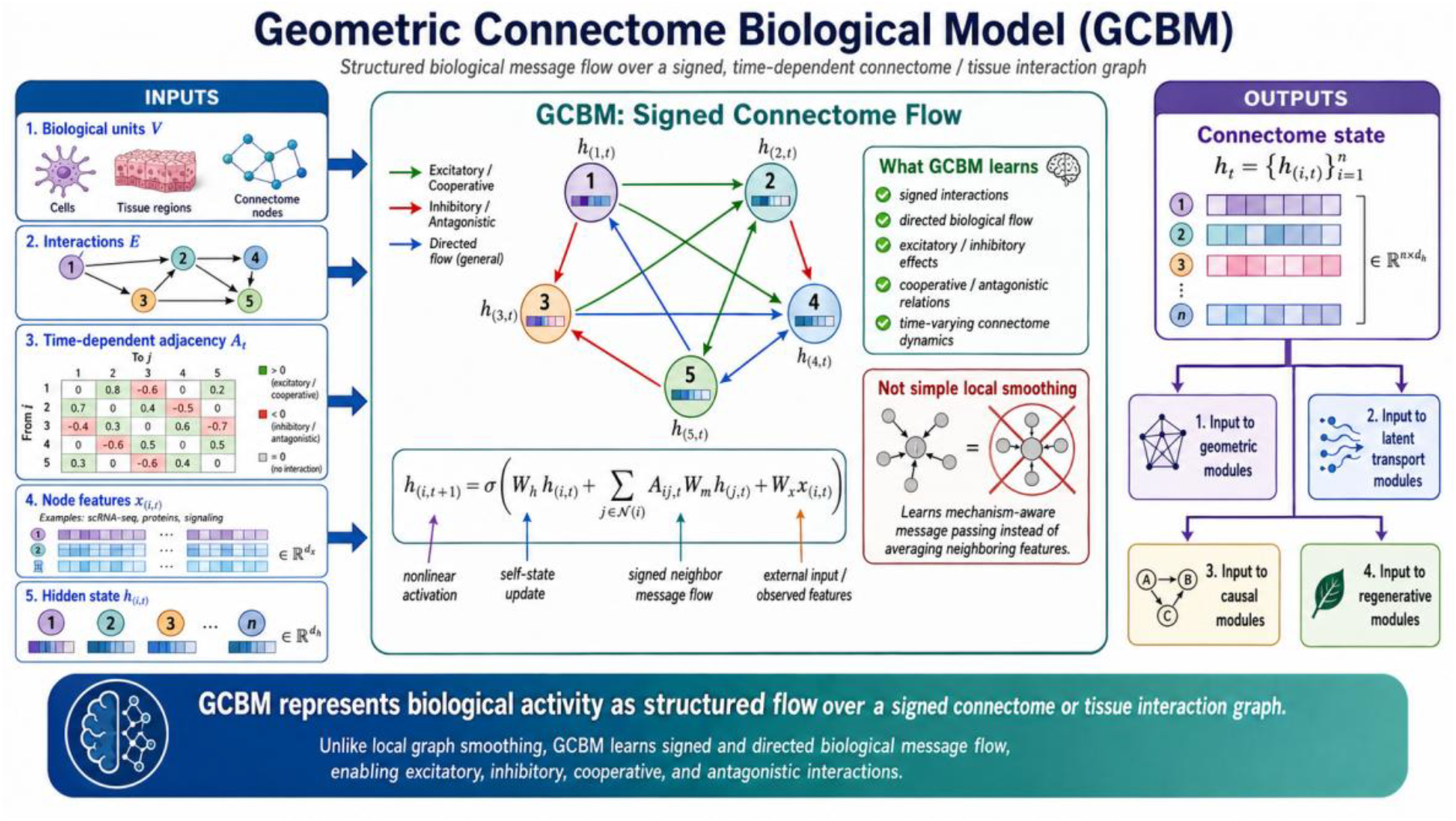
GCBM: Geometric Connectome Biological Model.

### 3.3 DCT: Differentiable Charted Tissue Geometry

Biological tissue states may lie on curved manifolds rather than flat Euclidean spaces. The **DCT** component models tissue geometry using differentiable charts.

A chart operator maps the latent state to local manifold (Xiong 2026) coordinates:

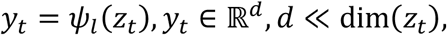

where *l* indexes a local chart. A decoder reconstructs the latent state:

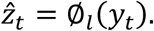

The geometry-preserving loss is

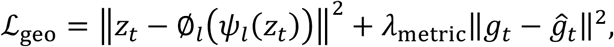

where *g*_*t*_ is the local metric tensor and 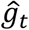 is the learned metric induced by the chart. For overlapping charts *l* and *k*, transition consistency is imposed by

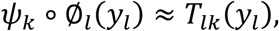

where *T*_*lk*_ is a learned transition map. This allows the model to represent local tissue geometry while maintaining global coherence.

DCT is novel relative to the baselines because it represents biological state as a charted manifold rather than as a flat vector or local graph average.

### 3.4 HV: Hamiltonian Latent Transport and Verification

The **HV** component introduces a structured latent dynamical system. The latent biological state is decomposed as

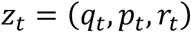

where *q*_*t*_ represents configuration-like biological state, *p*_*t*_ represents transformation momentum, and *r*_*t*_ represents mechanism-level latent coordinates.

A Hamiltonian function is learned:

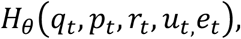

and the latent dynamics are modeled as

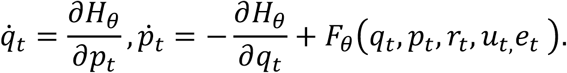

The additional force term *F*_*θ*_ allows non-conservative biological effects, such as growth, degradation, intervention response, and regeneration.

The dynamical loss is

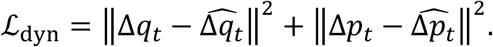

The verification component evaluates whether predicted transitions remain biologically and geometrically plausible:

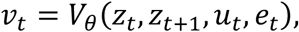

which define the verifier as a **probabilistic validity classifier** that takes the current state, predicted next state, intervention, and environment/domain context, then outputs a scalar plausibility score and a useful explicit form is:

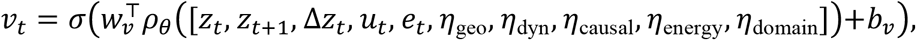

where:

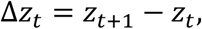

*ρ*_*θ*_(⋅)is a neural verifier encoder (Zhu et al. 2026), *w*_*v*_and *b*_*v*_are classifier parameters, and *σ*(⋅)is the sigmoid function:

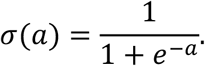

So *v*_*t*_ *≈* 1means the transition is judged biologically/geometrically valid, while *v*_*t*_ *≈* 0means it is judged implausible.

The verification loss is

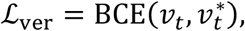

where 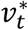 denotes a validity label or surrogate consistency target and the binary cross-entropy loss is:

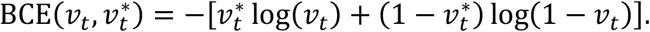

For numerical stability, use:

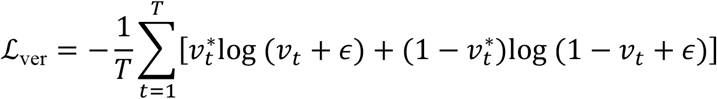

where *ϵ* > 0, for example:

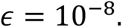

If a transition is experimentally observed and biologically valid:

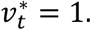

If it is known to be invalid, impossible, toxic, or biologically inconsistent:

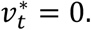

HV differs from the baselines because it models transformation direction, dynamical consistency, and biological plausibility, rather than only static prediction (Figure 3).

**Figure 3.**
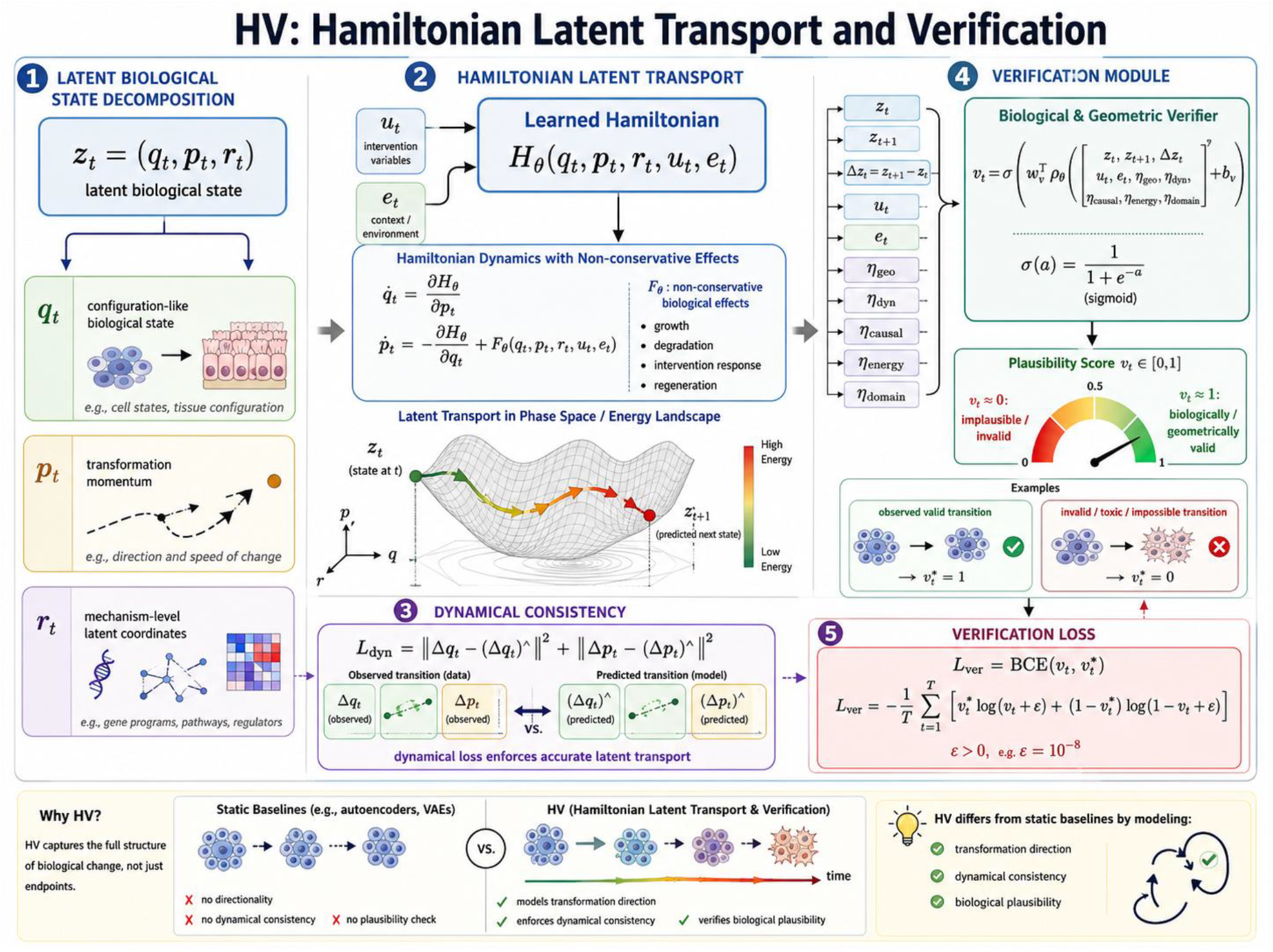
HV: Hamiltonian Latent Transport and Verification.

### 3.5 Bio: Nonlinear Biological Mechanism Layer

The **Bio** component explicitly represents nonlinear biological mechanism. The biological response is modeled as a coupled mechano-chemical-morphological process.

Here is a clean explicit mathematical formulation for the biological output function and interaction terms.

Let the biological latent/context variables be

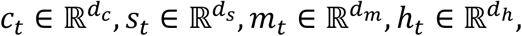

where, for example, *c*_*t*_ may represent cellular state, *s*_*t*_ spatial or tissue state, *m*_*t*_ molecular/mechanistic state, and *h*_*t*_ hidden regulatory context. Let *u*_*t*_be an intervention/control variable and *e*_*t*_ an environment/domain variable.

The biological prediction module is

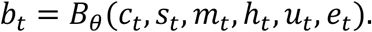

A useful explicit form is to define an interaction-augmented feature vector

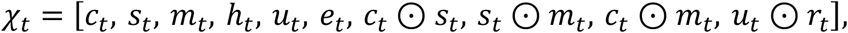

where ⊙denotes elementwise Hadamard multiplication.

Then the biological output can be modeled as

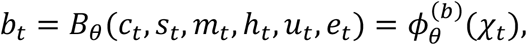

where 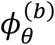 is a neural biological decoder or predictor.

A simple multilayer explicit form is

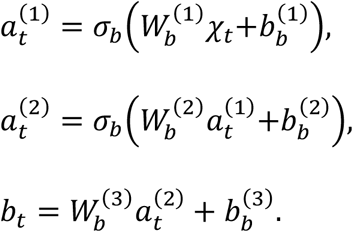

If *b*_*t*_represents bounded biological quantities, such as probabilities, expression proportions, viability scores, or normalized phenotype values, then one may use

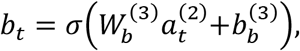

where

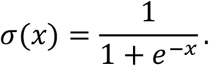

The interaction terms are explicitly

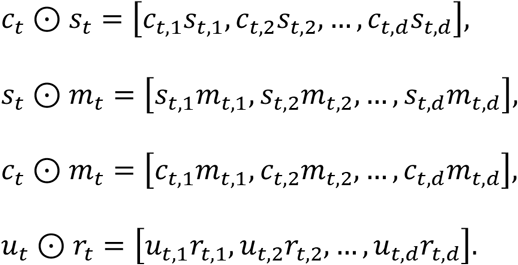

If the dimensions differ, each variable can first be projected into a shared interaction space:

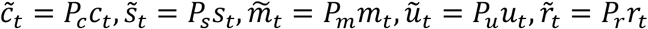

Then the interaction vector becomes

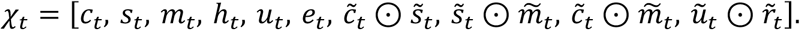

The biological reconstruction or prediction loss is

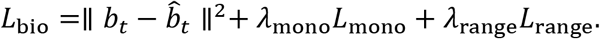

Here 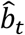 denotes the observed or target biological output.

A useful monotonicity penalty is

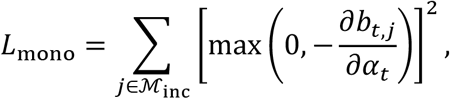

where α_*t*_ is a biological progression, dose, time, injury, regeneration, or intervention-intensity variable, and ℳ indexes outputs expected to increase monotonically.

For outputs expected to decrease monotonically, use

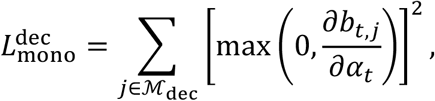

A combined monotonicity loss is therefore

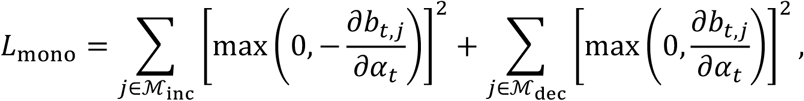

A useful range constraint is

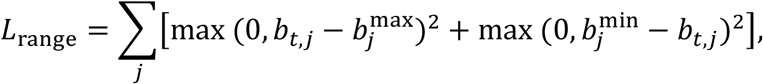

which penalizes predicted biological quantities that leave their admissible biological range

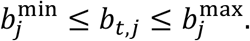

Thus the full explicit objective is

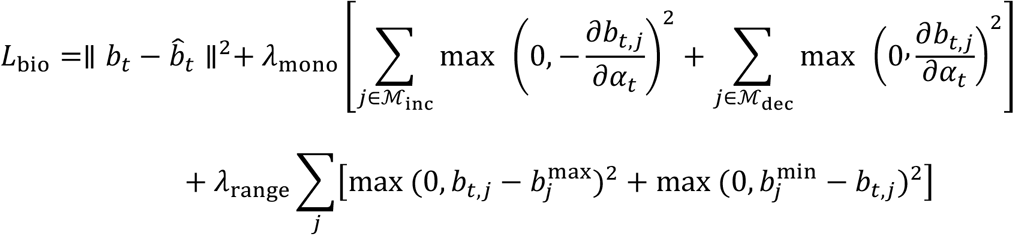

This formulation makes *B*_*θ*_explicitly depend not only on individual biological variables, but also on pairwise biological interactions such as cell–space, space–mechanism, cell–mechanism, and intervention–mechanism coupling (Figure 4).

**Figure 4.**
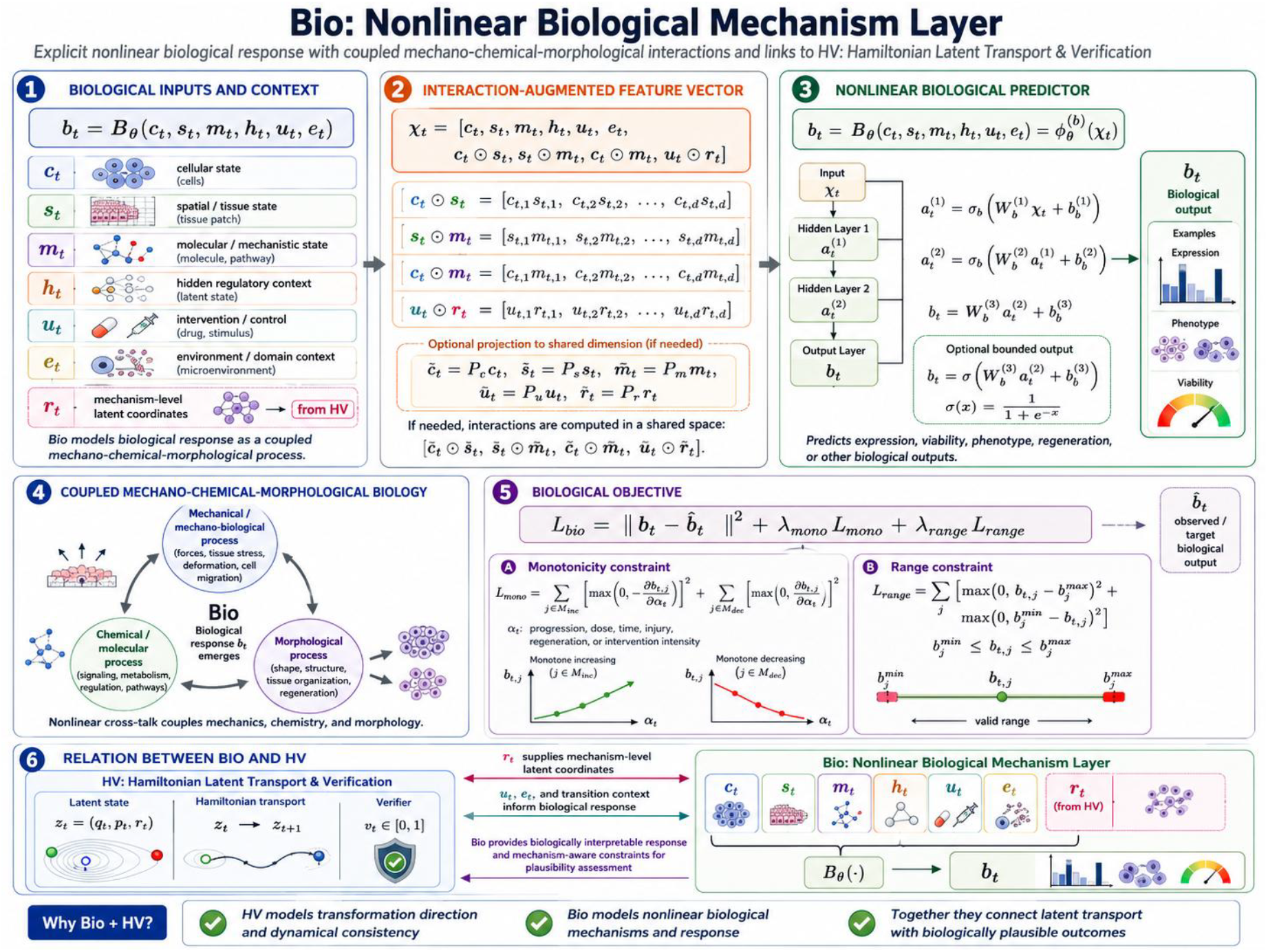
Bio: Nonlinear Biological Mechanism Layer.

### 3.6 NL: Nested Latent and Nested Memory Structure

The **NL** component represents biological information using nested latent memory channels. Instead of a single undifferentiated memory vector, the model separates memory into multiple functional channels:

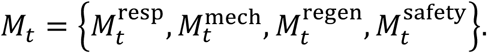

These channels represent response memory, mechanism memory, regeneration memory, and safety or validity memory.

For a current latent state *z*_*t*_, memory retrieval is defined by

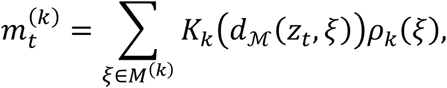

where *k* indexes the memory channel, *d*_ℳ_ is geodesic distance on the learned manifold, *K*_*k*_ is a kernel, and*ρ*_*k*_ is a memory density.

A task-aware gate combines memory channels:

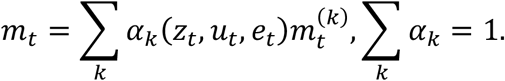

This nested memory structure differs from both baselines because it retrieves prior biological experience according to mechanism, regeneration, response, and safety relevance rather than simply using nearby samples.

### 3.7 Grow: Continual Growth Without Overwriting

The **Grow** component allows the model to incorporate new biological regimes without overwriting previously learned mechanisms. When the model encounters a new regime, it can allocate new parameters or memory slots:

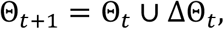

where ΔΘ_*t*_ is activated only when novelty or mechanism uncertainty exceeds a threshold. A novelty score is defined as

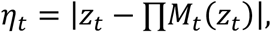

where ∏*M*_*t*_ projects the current state onto the previously learned manifold. If *η*_*t*_ is large, the model grows new local capacity.

The growth loss is

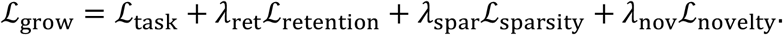

The retention term prevents catastrophic forgetting :

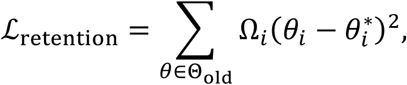

where Ω_*i*_ measures parameter importance.

Grow is novel relative to the baselines because it allows the model to expand under new biological regimes while preserving old knowledge.

### 3.8 CHG: Causal Hypergraph Structure

The **CHG** component models higher-order causal interactions. A biological response may not be caused by a single variable or pairwise edge, but by a combination of chemical, mechanical, morphological, and intervention variables.

A causal hypergraph is defined as

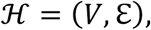

where each hyperedge

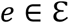

connects a set of variables

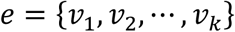

to a response or mechanism.

The hypergraph message is

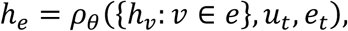

and node updates are

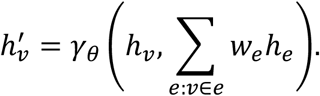

The hyperedge weight *w*_*e*_ represents the strength of a higher-order causal interaction.

CHG differs from local graph models because it captures multi-variable causal mechanisms rather than only pairwise local neighborhoods (Figure 5).

**Figure 5.**
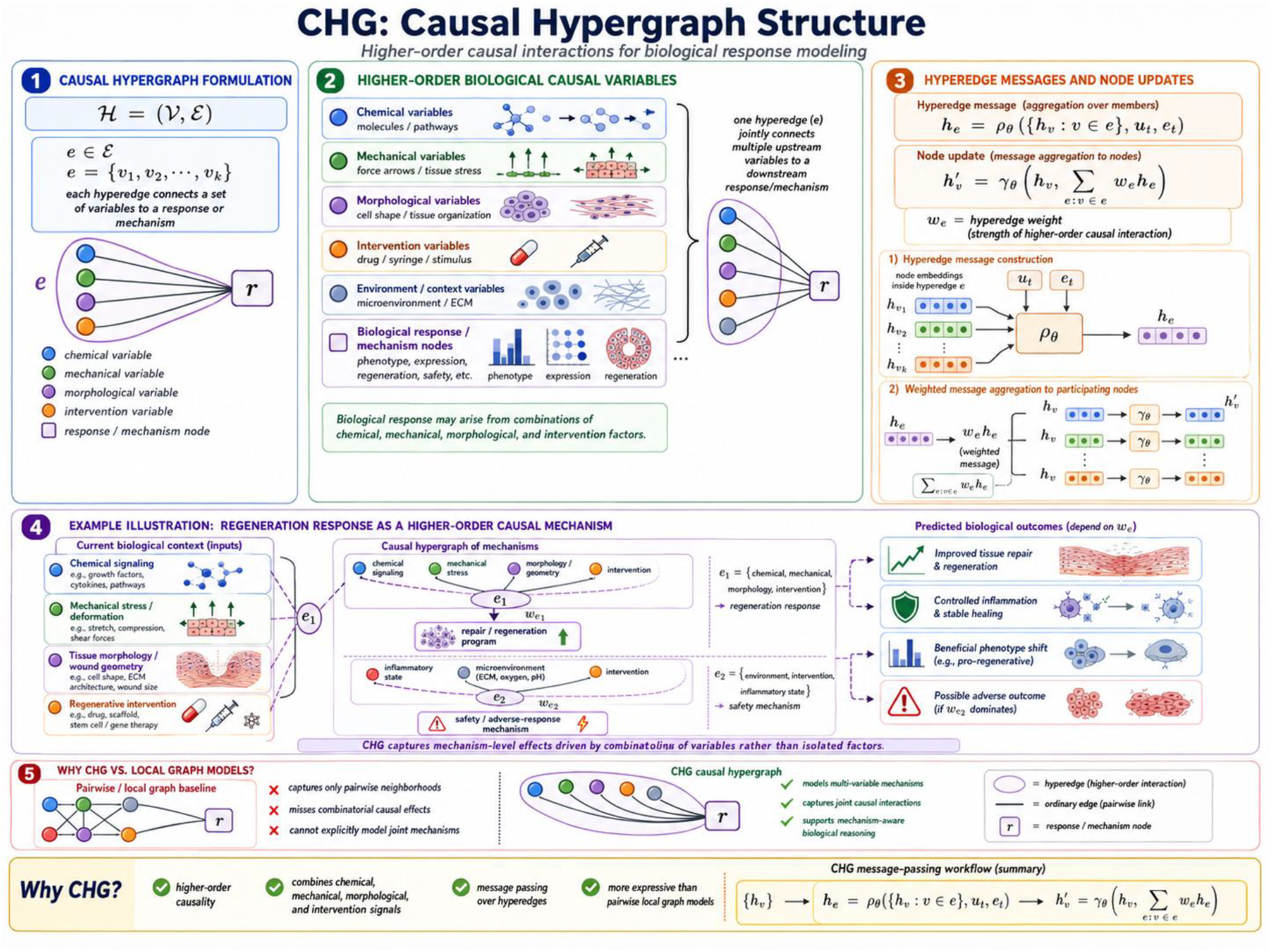
CHG: Causal Hypergraph Structure.

### 3.9 CSM: Causal Structure Module

The **CSM** component learns a mechanism-level causal structure:

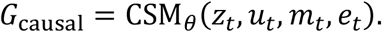

The CSM outputs a causal adjacency or hyper-adjacency structure that determines which latent mechanisms are active under a given context.

A structural causal representation is defined by

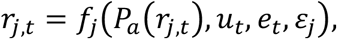

where *P*_*a*_(*r*_*j,t*_) denotes causal parents of mechanism coordinate *r*_*j*_.

The causal loss combines intervention prediction and sparsity:

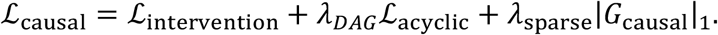

The acyclicity penalty may be written as

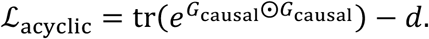

The intervention prediction loss is

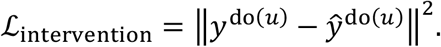

CSM is a key novelty because it separates causal mechanism discovery from ordinary predictive correlation.

### 3.10 RHEC: Regenerative Homeostatic Error Correction

The **RHEC** component models biological correction toward viable, regenerative, or homeostatic states. Many tissue systems respond to perturbation by attempting to restore structure, repair damage, or move toward stable attractors.

Let *z*_*t*_ be the predicted biological latent state and 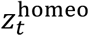 be a learned homeostatic reference state. The homeostatic error is

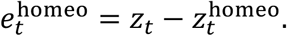

The regenerative correction field is

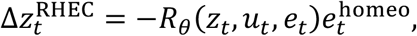

where *R*_*θ*_(*z*_*t*_, *u*_*t*_, *e*_*t*_)is a learned correction operator.

The corrected latent state is

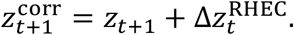

The RHEC loss is

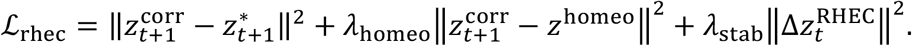

RHEC is especially important under regeneration and out-of-distribution shifts. It differs from Euclidean and local graph baselines because it explicitly models correction toward biologically plausible attractor states.

### 3.11 Full Prediction Model

The full model combines all components as follows (Figure 6):

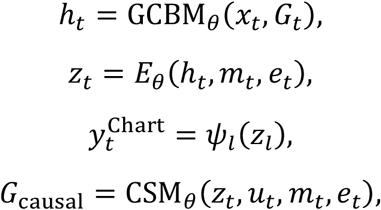

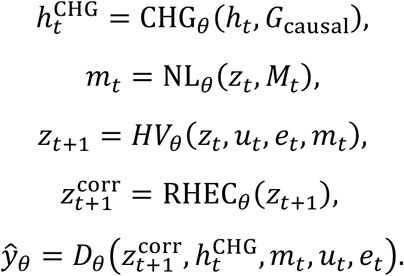

**Figure 6.**
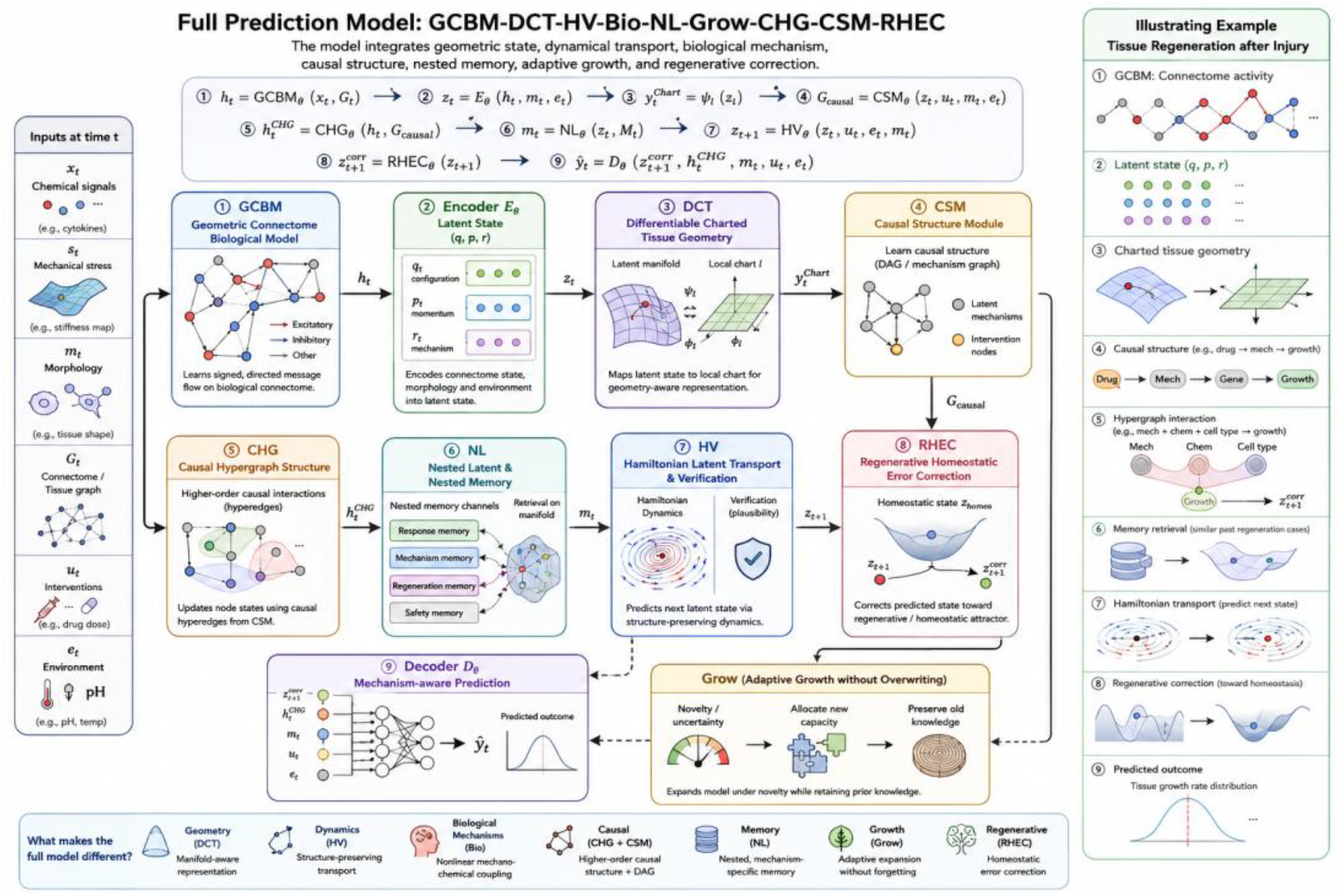
Full Prediction Model: GCBM-DCT-HV-Bio-NL-Grow-CHG-CSM-RHEC.

The model therefore predicts outcomes using geometric state, dynamical transport, biological mechanism, causal structure, nested memory, adaptive growth, and regenerative correction.

## 4. Synthetic Toy Studies

We evaluated the full model using four synthetic toy studies designed to test progressively more difficult biological prediction settings.

### 4.1 Toy A: Local Mechano-Chemical Regression

Toy A represents a relatively simple setting in which the response depends on local chemical signal, mechanical stiffness, and tissue morphology. This setting contains smooth local structure and moderate nonlinear interaction.

### 4.2 Toy B: Nonlinear Biological Mechanism

Toy B introduces stronger nonlinear coupling among chemical signal, mechanical stress, and morphology. The purpose is to test whether the model can capture nonlinear biological mechanisms that are not well represented by Euclidean or local graph baselines.

### 4.3 Toy C: Causal Intervention Response

Toy C introduces intervention variables and mechanism-dependent response. The purpose is to test whether causal and hypergraph components improve prediction when the response depends on intervention-sensitive latent mechanisms.

### 4.4 Toy D: Regenerative Out-of-Distribution Shift

Toy D introduces an out-of-distribution shift and regenerative response. This setting tests whether the model can remain accurate when local graph similarity becomes unreliable and when the system moves toward a new homeostatic or regenerative regime.

## 5. Evaluation Metrics

The primary metric is mean squared error:

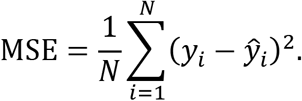

We also report mean information gain relative to the Euclidean baseline:

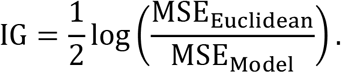

Euclidean information gain is zero by definition. Positive values indicate improvement over the Euclidean baseline.

## 6. Results

Across 50 repeated synthetic simulations, the full GCBM-DCT-HV-Bio-NL-Grow-CHG-CSM-RHEC model achieved the lowest MSE on all four toy studies.

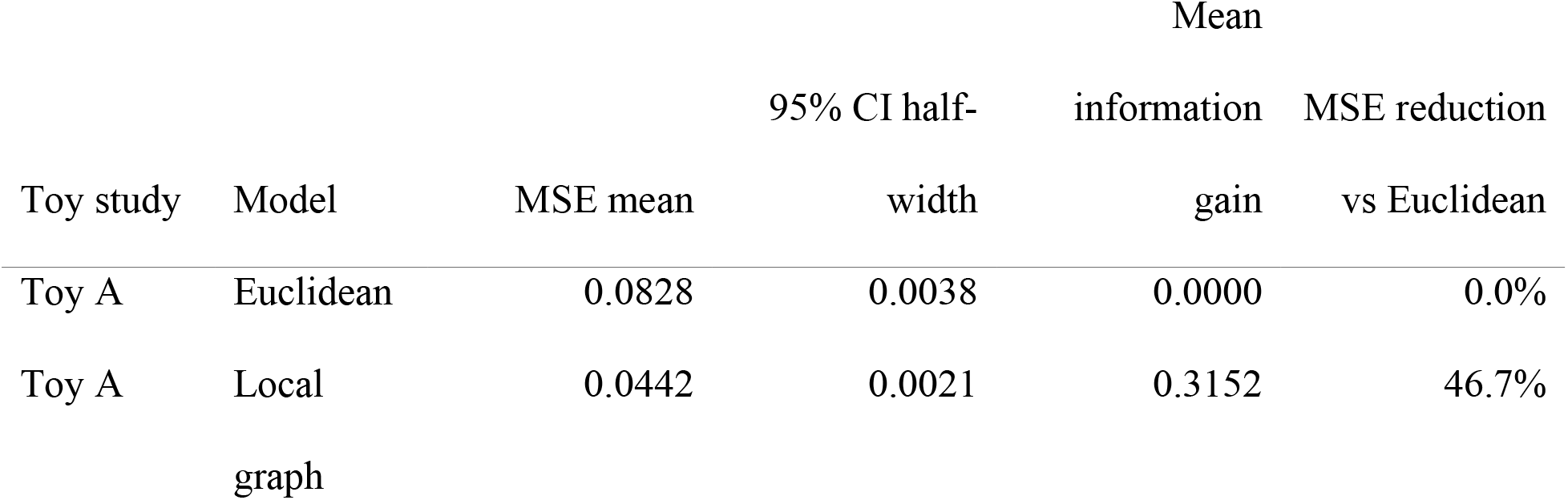
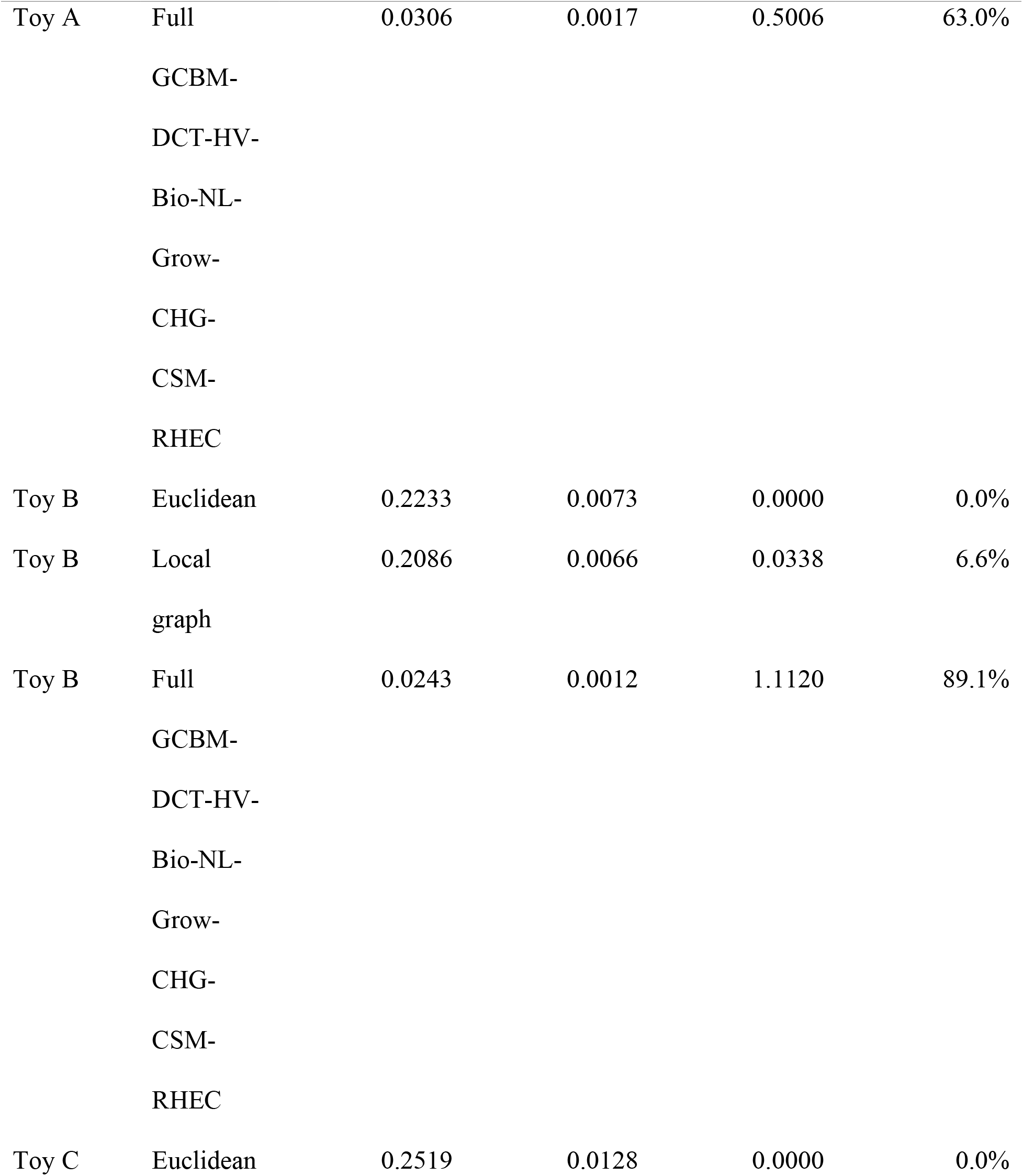
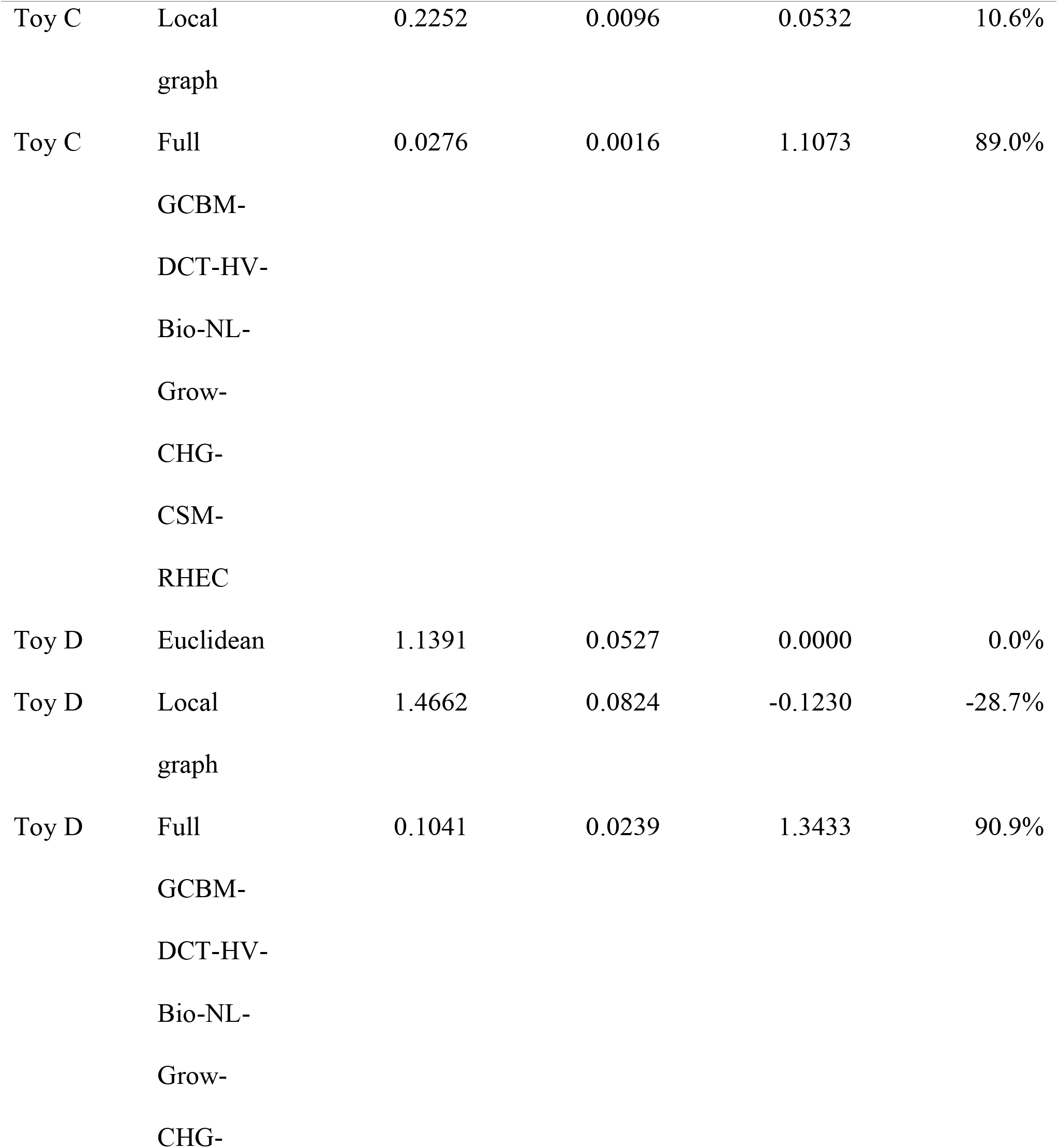
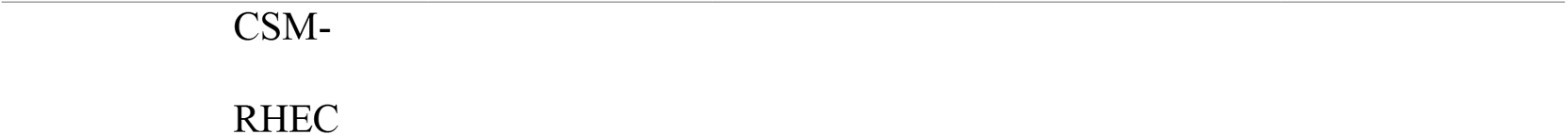

The local graph baseline improved over Euclidean in Toy A, Toy B, and Toy C, but the magnitude of improvement was small in the nonlinear and causal settings. In Toy D, the local graph baseline performed worse than the Euclidean baseline, suggesting that local smoothing can be harmful under out-of-distribution regenerative shifts. In contrast, the full model remained robust and achieved the strongest performance in every toy setting.

## 7. Discussion

The results suggest that biological prediction benefits from models that explicitly encode geometry, mechanism, causal structure, and regenerative correction. The Euclidean baseline performed reasonably only when the response was dominated by raw covariates. The local graph baseline improved prediction when neighborhood similarity was informative, but it failed under the most difficult regenerative shift setting. This indicates that graph locality alone is insufficient for modeling intervention-sensitive biological systems.

The full GCBM-DCT-HV-Bio-NL-Grow-CHG-CSM-RHEC model achieved the strongest performance because it combines multiple complementary inductive biases. GCBM captures signed connectome flow. DCT represents tissue state on differentiable local charts. HV models directionality and dynamical consistency. Bio captures nonlinear biological interactions. NL retrieves mechanism-specific memory. Grow allows adaptive expansion under novelty. CHG captures higher-order causal interactions. CSM estimates intervention-sensitive causal structure. RHEC corrects predictions toward regenerative and homeostatic plausibility.

These components are not independent add-ons; they form a unified architecture. The geometric modules identify the biological state space. The dynamical modules model movement through this space. The causal modules determine which mechanisms are active. The memory and growth modules preserve prior knowledge while adapting to new regimes. The regenerative correction module stabilizes predictions under biological repair or recovery.

The strongest advantage was observed in Toy D, where the local graph baseline failed. This result is important because biological systems frequently undergo shifts due to disease, injury, therapy, adaptation, or regeneration. A model that relies only on local similarity may propagate incorrect neighborhood information. The full model instead uses mechanism, causal structure, and homeostatic correction to maintain biological plausibility.

## 8. Limitations

The present study uses synthetic toy datasets rather than real biological datasets. Therefore, the numerical results should be interpreted as controlled simulation evidence rather than empirical validation on biological experiments. The model also contains many components, and future work should perform systematic ablation studies to quantify the contribution of each component. Additional work is needed to test scalability, identifiability of causal mechanisms, robustness to measurement noise, and interpretability of learned latent states.

## 9. Future Work

Future work should evaluate the model on real datasets involving single-cell perturbation response (Heidari et al. 2026), spatial transcriptomics, tissue regeneration, organoid development, connectome activity, and drug-response prediction. A detailed ablation study should compare the full model against versions that remove DCT, HV, Bio, NL, Grow, CHG, CSM, or RHEC. Another important direction is to formalize the learned causal mechanisms and test whether the model can propose experimentally testable biological hypotheses.

## 10. Conclusion

We introduced GCBM-DCT-HV-Bio-NL-Grow-CHG-CSM-RHEC, a unified framework for geometric, biological, causal, and regenerative modeling. The model differs fundamentally from Euclidean and local graph baselines by representing biological systems as structured dynamical objects on learned tissue manifolds with causal mechanisms, nested memory, adaptive growth, and homeostatic correction. In synthetic Toy A–D studies, the full model achieved the lowest MSE and highest mean information gain across all settings. These findings support the proposed architecture as a promising direction for mechanism-aware prediction in complex biological systems.

## Acknowledgements

The authors wish to acknowledge the use of AI-powered language models (ChatGPT) for assistance in creating images, improving the grammar, spelling, and readability of this manuscript.

## Author contributions

TX and ZH: Derive formulas and perform data analysis, XS: Problem formulation formula derivation, LJ: Design project, MX: Design project, perform data analysis and write paper.

